# Virological characteristics of SARS-CoV-2 Omicron BA.5.2.48

**DOI:** 10.1101/2024.03.26.586802

**Authors:** Wenqi Wang, Qiushi Jin, Ruixue Liu, Wentao Zeng, Pengfei Zhu, Tingting Li, Tiecheng Wang, Haiyang Xiang, Hang Zhang, Qin Chen, Yun Gao, Yana Lai, Fang Yan, Xianzhu Xia, Jianmin Li, Xuefeng Wang, Yuwei Gao

## Abstract

With the prevalence of sequentially-emerged sublineages including BA.1, BA.2 and BA.5, SARS-CoV-2 Omicron infection has transformed into a regional epidemic disease. As a sublineage of BA.5, the BA.5.2.48 outbreak and evolved into multi-subvariants in China without clearly established virological characteristics, especially the pathogenicity. Though reduced airborne transmission and pathogenicity of former Omicron sublineages have been revealed in animal models, the virological characteristics of BA.5.2.48 was unidentified. Here, we evaluated the in vitro and in vivo virological characteristics of two isolates of the prevalent BA.5.2.48 subvariant, DY.2 and DY.1.1 (a subvariant of DY.1). DY.2 replicates more efficiently than DY.1.1 in Hela^hACE2+^ cells and Calu-3 cells. The A570S mutation (of DY.1) in a normal BA.5 spike protein (DY.2) leads to a 20% improvement in the hACE2 binding affinity, which is slightly reduced by a further K147E mutation (of DY.1.1). Compared to the normal BA.5 spike, the double-mutated protein demonstrates efficient cleavage and reduced fusogenicity. BA.5.2.48 demonstrated enhanced airborne transmission capacity in hamsters than BA.2. The pathogenicity of BA.5.2.48 is greater than BA.2, as revealed in K18-hACE2 rodents. Under immune selection pressure, DY.1.1 shows stronger fitness than DY.2 in hamster turbinates. Thus the outbreaking prevalent BA.5.2.48 multisubvariants exhibites divergent virological features.

**Importance:** Omicron continues to circulate and evolves novel sublineages with indistinguishable pathogenicity and transmission. Therefore humanized Omicron-sensitive animal models must be applied to evaluate the virological characteritics and antiviral therapeutics. By using multiple models including the Omicron-lethal H11-K18-hACE2 rodents, BA.5.2.48 revealed higher pathogenicity in the novel H11-K18-hACE2 rodent models than the previously epidemic BA.2, and thus the models are more adapted to Omicron studies. Moreover, the regional outbreaking of BA.5.2.48 promotes the multidirectional evolution of its subvariants, gaining either enhanced pathogenicity or a fitness in upper airways which is associated with higher transmission, highlighting the importance of surveillance and virological studies on regionally endemic sublineages which represents the short-run evolutionary direction of Omicron.

## Introduction

The Omicron variant of severe acute respiratory syndrome coronavirus 2 (SARS-CoV-2) emerged at the end of 2021 and has evolved into complex sublineages. Until October 2022, three major Omicron lineages had serially transitioned into globally dominant variants of concern (VOCs): first, BA.1, followed by BA.2, and then its sublineage BA.5. BA.5, along with its subvariants BA.5.1 and BA.5.2, were the predominant strains worldwide for several months, during which the spike protein of these strains was found to exhibit greater hACE2 binding affinity than those of BA.1 and BA.2(1); further, the BA.5 strains demonstrated antibody evasion(2). Moreover, BA.5 has been revealed to have similar or slightly enhanced pathogenicity than BA.2 in rodent models(3–5).

Before November 2022, China employed a precise prevention and control strategy (the dynamic zero-COVID-19 policy) to control regional outbreak clusters of COVID-19; however, this policy ended in mid-November 2022. Subsequently, the number of COVID-19 cases was estimated to reach hundreds of millions(6), of which one of the most predominant sublineages was BA.5.2.48 (and its subvariants)(7, 8). Thus, the Omicron BA.5.2.48 epidemic in China prompted evolutionary divergence, resulting in genomic multiplicity of the virus, of which novel mutations introducing virological characteristics triggered concern. The BA.5.2.48 in China include DY.1 (Alias of BA.5.2.48.1), DY.2, DY.3 and DY.4 sublineages; only DY.1 bears a spike protein mutation (A570S), and its subvariant DY.1.1 bears an additional mutation, namely, K147E; both are expected to contribute to antigenicity shifts and alterations in virological characteristics. Moreover, DY.2 and DY.1 became the most predominant subvariants of BA.5 in China in the period before XBB subvariants prevailed (see Results for details), though their characteristics were unidentified.

Here, the in vitro replicative kinetics of the isolates of DY.1.1 and DY.2 subvariants were compared with those of the former epidemic Omicron BA.1 and BA.2. Further, the pathogenicity of DY.1.1 and DY.2 in wild-type (WT) and K18-hACE2 rodents was revealed. We also demonstrated the in vivo comparative fitness of DY.1.1 and DY.2 under immune pressure. Moreover, as the spike universally impacts infection and pathogenicity of SARS-CoV-2 via its functional characteristics(9–12), the changes in spike features were studied to explain the differences in virological characteristics among the Omicron strains studied here.

## Results

### Prevalence and mutations of BA.5.2.48 and its subvariants in China

On the basis of the BA.5 genome, the BA.5.2 variant bears the synonymous mutation A28330G; a variant with a further T1050N amino acid mutation in the ORF1b protein (Fig. 1A) was defined as BA.5.2.48. As a subvariant of BA.5.2.48, DY.1 bears the A570S mutation in the spike protein, and DY.1.1 bears three additional amino acid mutations (including K147E in the spike). DY.2, another BA.5.2.48 subvariant, bears a Q241K mutation in the nucleocapsid protein. Moreover, as another prevailing subvariant of BA.5.2, BA.5.2.49 bears an amino acid mutation in the spike (T883I). From the end of the dynamic zero-COVID-19 policy in mid-November 2022 to April 2023 when the XBB subvariant predominated, BA.5.2.48 and BF.7.14 were the two most prevalent sublineages in China during this period. Among the four BA.5.2.48 sublineages (DY.1, DY.2, DY.3 and DY.4), only the prevalence of DY.1 and DY.2 was demonstrated to keep increasing after emergence, and accounted for up to 11% and 18% of new infections, respectively (Fig. 1B). Thus, here, we investigated the virological characteristics of DY.1 and DY.2.

**Fig. 1.**
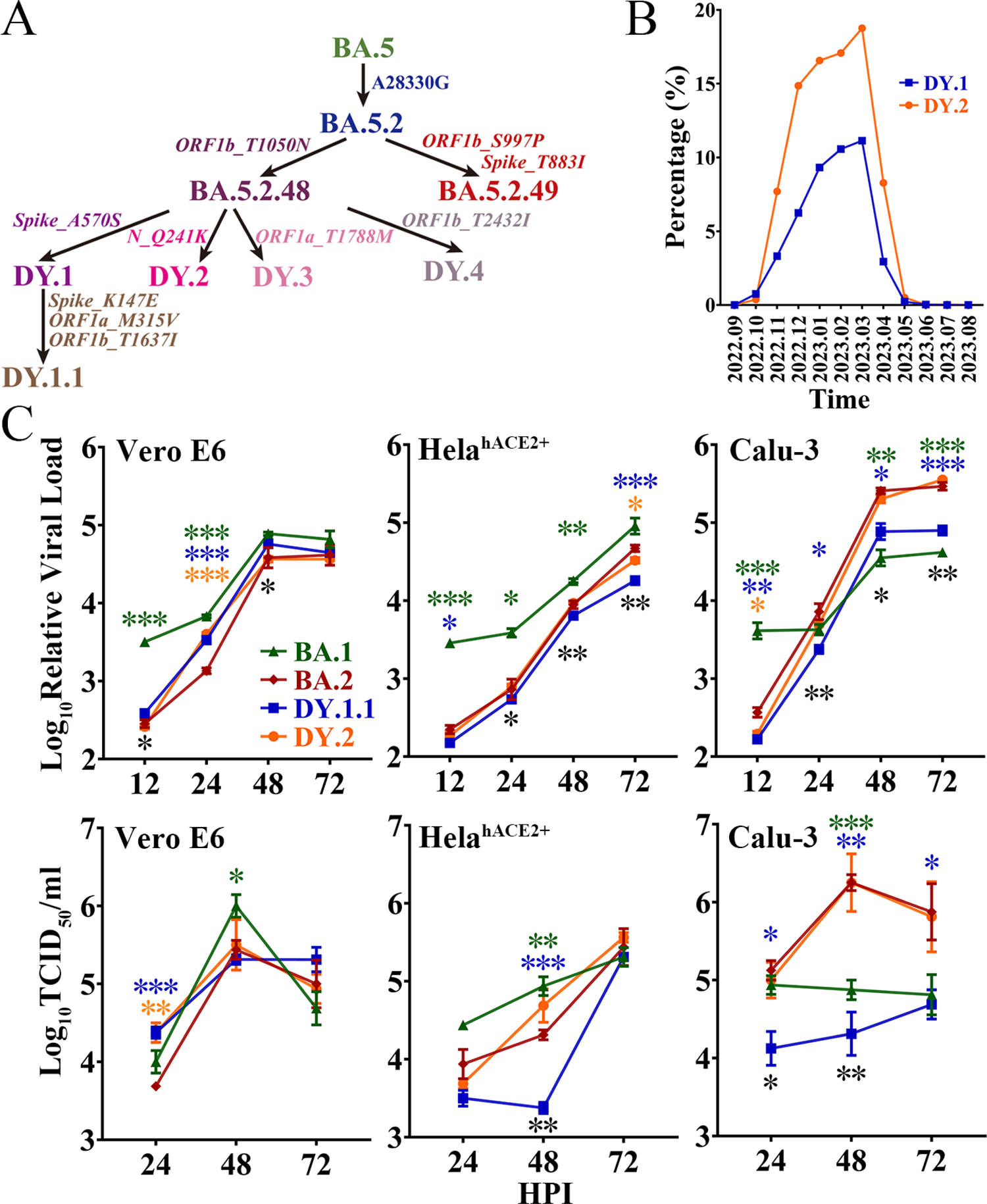
Evolution, prevalence and replicative kinetics of DY.1 and DY.2. **(A)** Evolutionary origin of BA.5.2.48 sublineages, including DY.1 and its sublineages DY.1.1, DY.2, DY.3 and DY.4. Synonymous mutations in nucleotides and amino acid mutations are shown in regular and italic font, respectively. **(B)** Prevalence of DY.1 (blue) and DY.2 (orange) in China for one year from September 2022 (2022.09) to August 2023 (2023.08). **(C)** Replicative kinetics of DY.1.1 and DY.2. The viral loads (upper panel) and viral titers (lower panel) of the Omicron BA.1, BA.2 and BA.5.2.48 isolates were determined in Vero E6 cells, HeLa^hACE2+^ cells and Calu-3 cells. The significance of the differences in replication between BA.2 (dark red) and BA.1 (green), DY.1.1 (blue) or DY.2-CC1 (orange) are indicated above the lines by the asterisks in colors corresponding to the individual virus. The significance of the differences in replication between DY.1.1 and DY.2 is indicated by the black asterisks below the lines.

### In vitro replication of Omicron BA.1, BA.2 and BA.5.2.48

The in vitro replication of BA.1, BA.2 and two BA.5.2.48 isolates (DY.1.1 and DY.2) was evaluated in three cell lines, namely, Vero E6 cells, which are commonly used for SARS-CoV-2 isolation and passage; HeLa cells overexpressing hACE2 (HeLa^hACE2+^ cells); and human lung Calu-3 cells. Because of differences in origin, passaging and culture conditions, the same cell type could have distinct gene expression patterns that strikingly affect viral replication. RNA sequencing analysis of the three cell types in our laboratory was performed to evaluate the expression of key SARS-CoV-2 entry-related genes, including the major acceptor ACE2 and the serine protease TMPRSS2. Both Vero E6 cells and HeLa^hACE2+^ cells were TMPRSS2 negative, although they expressed ACE2 at quite low and exceedingly high levels, respectively (Table S3). Calu-3 cells expressed a moderate level of ACE2 of which TMPRSS2 was positive.

The replication kinetics were first characterized according to the viral load. Reproduced in Vero E6 cells, DY.1.1 presented higher viral loads than DY.2 at 12 hours post infection (HPI) and 48 HPI (Fig. 1C). No significant differences in final viral loads were detected among the Omicron isolates, although BA.1, DY.1.1 and DY.2 replicated faster than BA.2. However, in both HeLe^hACE2+^ cells and Calu-3 cells, DY.1.1 presented a consistently lower viral load than DY.2. Moreover, very few discrepancies in replicative kinetics were demonstrated between DY.2 and BA.2 in all the three cell types. The replication discrepancies in various Omicron viruses revealed by viral titer generally resembled those revealed by viral load. There were no significant differences among the final viral titers of BA.1, BA.2 and the two BA.5.2.48 isolates in Vero E6 cells and HeLe^hACE2+^ cells, although BA.1 replicated faster and DY.1.1 replicated slower. However, in Calu-3 cells, DY.1.1 and BA.1 exhibited weak propagation, which suggest that the Omicron strains shows differences in TMPRSS2 usage to entry cells. In addition, all Omicron strains demonstrated a reduced replication than WT in Vero E6 cells (Fig. S1). Furthermore, as indicated by both the viral load and titer, replication discrepancies between the two DY.2 isolates (CC1 and CC2) were hardly detectable (Fig. S2), although genomic differences did exist between them. Thus, only DY.2-CC1 was used for further studies.

### Characteristics of BA.5.2.48 spike mutations

To investigate the mutation-induced characteristics of the BA.5.2.48 spike, spike-mediated infectivity was firstly determined by pseudovirus-based infection. The A570S mutation (of DY.1) significantly reduced the infectivity of the normal BA.5 spike in hACE2-expressing cells, which was interestingly recovered by a further K147E mutation (of DY.1.1, Fig. 2A). Significantly, both the normal BA.5 spike and the mutants were associated with stronger infection than BA.2 spike (p<0.05); furthermore, the D614G spike was associated with greater infection than the spikes of BA.2 and BA.5 (p<0.05).

**Fig. 2.**
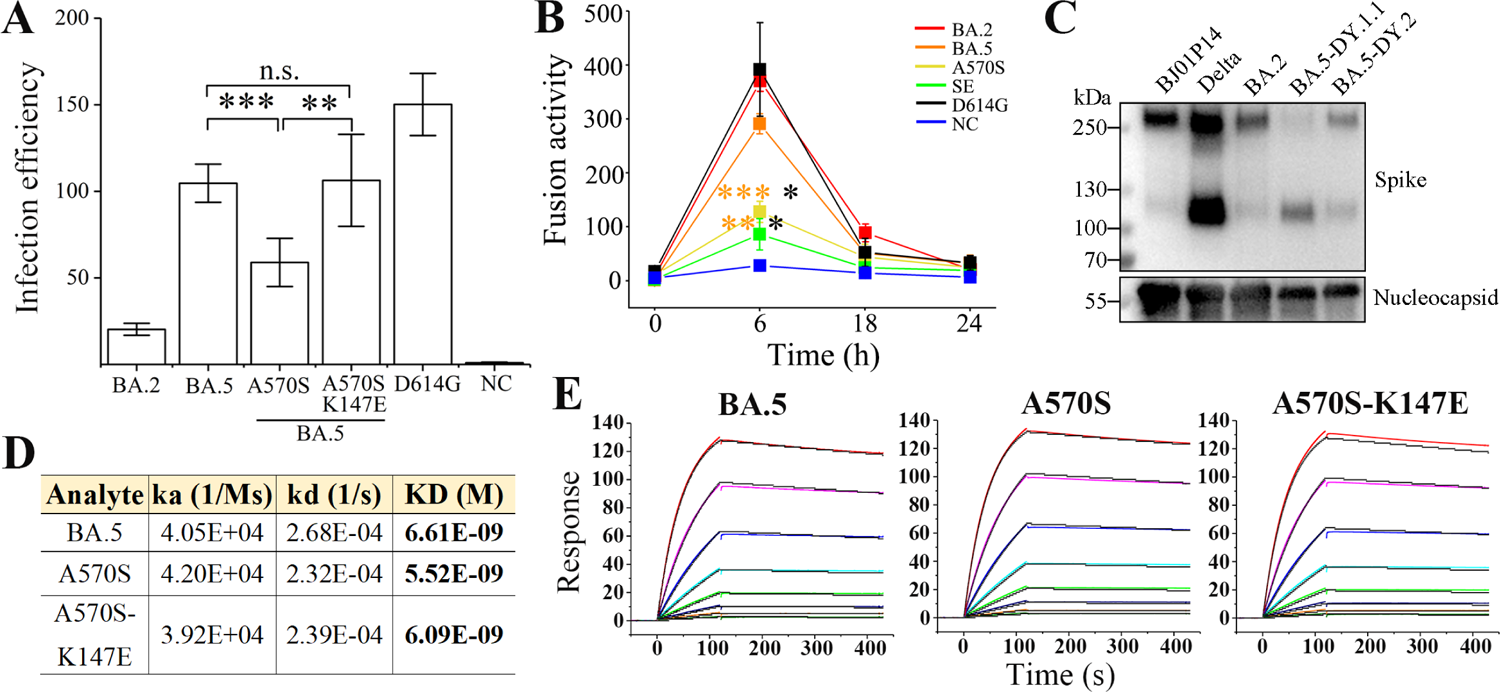
Characteristics of the spikes of BA.5.2.48 subvariants. **(A)** Spike-mediated infection by pseudovirus assay. The infectivity of the strains relative to that of the negative control (NC) is shown. **(B)** Spike-mediated cell-cell fusion determined by the DSP method. Significant differences between the spikes of previous epidemic variants (including D614G in black, BA.2 in red and BA.5 in orange) and the spikes of BA.5-A570S (A570S, yellow) or BA.5-A570S-K147E (SE, green) are revealed above the lines with asterisks in colors corresponding to the individual variants. **(C)** Spike procession in authentic SARS-CoV-2 virions, including BJ05P14, Delta, BA.2 and the two BA.5 subvariants DY.1.1 and DY.2. **(D-E)** Comparative binding affinities, including data (D) and response curves (E), of the BA.5, BA.5-A570S and BA.5-A570S-K147E spike to hACE2.

The formation of syncytia is regarded as a hallmark of SARS-CoV-2-induced pathogenesis in the lungs(13, 14) and is caused by spike-ACE2 interaction on the cell surface, referred to as spike-mediated cell-cell fusion(15). We first investigated the fusogenicity of BA.5 spikes using a method based on DSP. At 6 h post cell contact, a significant decrease in the fusogenicity induced by the A570S mutation was observed compared to that for the normal BA.5 spike, and the average fusogenicity was further reduced by the K147E mutation although no significant difference was demonstrated between the two mutants. (Figs. 2B and S3). Meanwhile, the fusogenicity of all the three BA.5 spikes was also significantly lower than that of BA.2 (p<0.01). However, despite the nonsignificant difference, the D614G were associated with a greater average fusogenicity than the BA.2 and BA.5 spikes. HeLa^hACE2+^ cells expressing the D614G spike also exhibited larger syncytia than the Omicron spikes (Fig. S3), which was similar to the GFP fluorescence results obtained via the DSP method (Fig. S4).

The activation of spike protein processing at the S1/S2 polybasic cleavage site on the virion is positively correlated with enhanced infection and fusogenicity(11, 12). To directly investigate the exact effect of the A570S/K147E double mutation on the cleavage of spike, we collected authentic SARS-CoV-2 virions and performed an immunoblot assay; the results revealed ∼250 kD bands corresponding to the full-length spike and ∼120 kD bands corresponding to the S2 subunit. The full-length band was hardly visible for the DY.1.1 spike with the A570S-K147E double mutation, while the normal BA.5 (DY.2) spike demonstrated approximately 20% cleavage (Fig. 2C). Spikes of both the DY.1.1 and DY.2 indicated stronger cleavage than the BA.2. Our results, consistent with those of a previous study(16), indicated that the Delta spike was strongly cleaved. Virions of another SARS-CoV-2 variant isolated in our laboratory (data not published, named BJ05P14) containing a deficient furin cleavage site in the spike barely exhibited spike cleavage. Interestingly, although the double mutation of the DY.1.1 spike induced improved processing, no enhanced spike-mediated infection or fusion was significantly demonstrated, which implies that the extent of spike processing is not directly related to the infection and fusogenicity it induces, at least for the BA.5 mutants studied here.

Omicron spikes demonstrated higher binding affinities than previous VOC(17). The binding affinities of the spike ectodomains of the BA.5.2.48 subvariants (Fig. S5) to hACE2 were measured via SPR (surface plasmon resonance). Compared to the normal BA.5 spike, the A570S mutant of DY.1 exhibited an approximately 20% enhanced binding affinity to hACE2, while the additional K147E mutation in the DY.1.1 spike slightly reduced the affinity (Fig. 2D and 2E). However, the A570S-K147E double mutant still demonstrated higher binding affinity than the BA.5 spike.

### Virological characteristics of DY.1.1 and DY.2 in vivo

The fast spread of BA.5.2.48 implies its probable high transmission capacity. Given that BA.5.2.48 spikes revealed highly improved processing which may induce efficient in vivo infection, the airborne transmission of BA.5.2.48 and BA.2 were compared in hamsters. Despite the successful inoculation of all donor hamsters, BA.2 was airborne transmitted at a frequency of 20% (1 of 5 animals). Moreover, we only detected a quiet inefficient infection (approximately 30 TCID_50_/ml) in the turbinate (other than the lung) of the transmitted animal. In the contrary, BA.5.2.48 (DY.1.1) was transmitted 60% (3 of 5 animals), and both lungs and turbinates were infected.

We evaluated the pathogenicity of BA.2, DY.1.1 and DY.2 in both WT and hACE2 transgenic Syrian hamsters. All the infected WT animals survived after challenge (Fig. 3B). Despite the significantly lower weights of the DY.1.1- and DY.2-infected hamsters than the mock-infected hamsters, most of the animals infected with the three Omicron isolates gained weight over the ten-day experiment, although weight loss was observed in a few BA.2-infected individuals (Fig. 3C). Moreover, the viral loads in the nasal lavages of each infection group continued to decrease from 2 to 8 DPI (days post infection) (Fig. 3D). Although the DY.1.1 viral loads were higher than those of BA.2 at 2 DPI, no significant differences were detected among the BA.2-, DY.1.1- and DY.2-infected groups from 4 to 8 DPI. In contrast, the BA.2-infected group demonstrated higher viral titers in turbinates than the DY.1.1-infected group at 3 DPI (Fig. 3E). No significant differences in viral titers were revealed in the lungs of the three groups. Divergent to nasal lavage and turbinate results, all three Omicron infections resulted in large intragroup differences in viral titers in the lower airways, which is highly consistent with previous results(3).

**Fig. 3.**
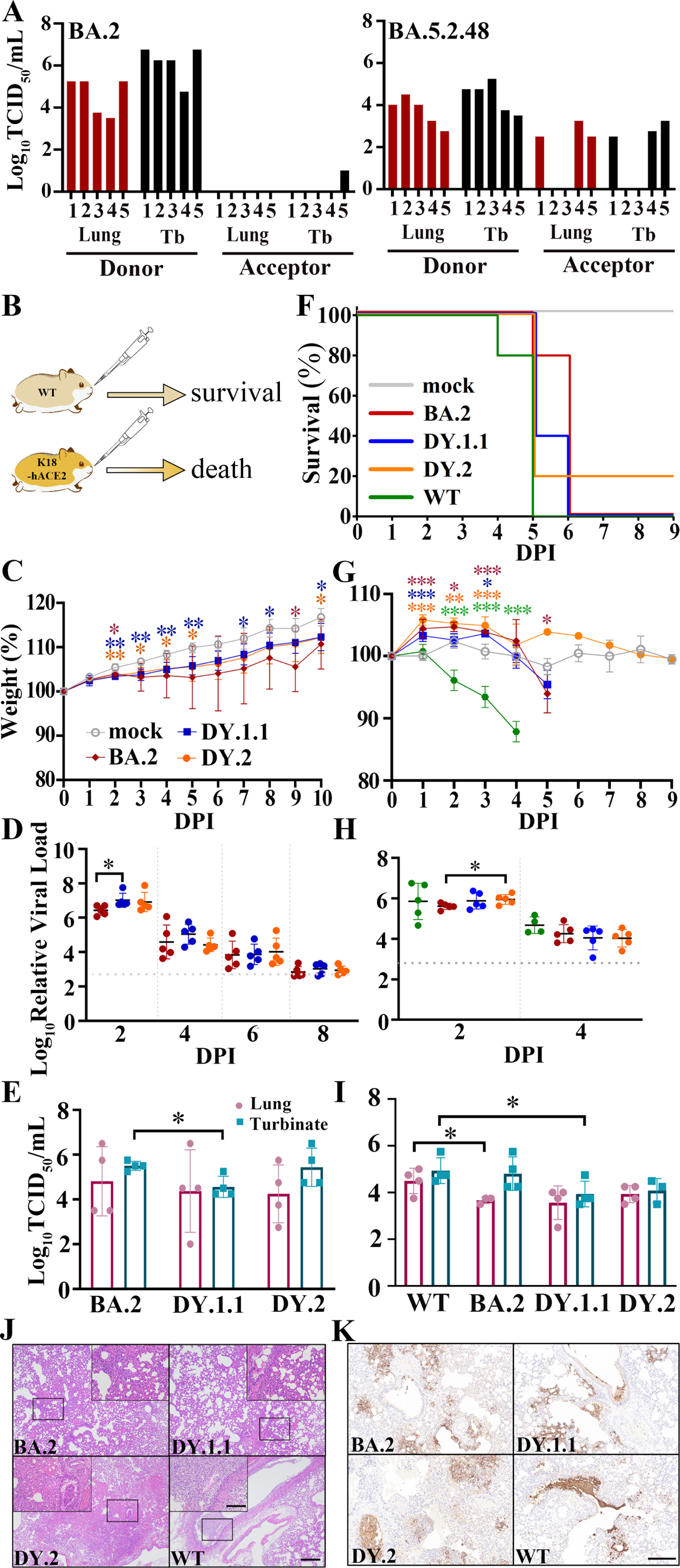
In vivo virological characteristics of BA.5.2.48 in hamsters. **(A)** Airborne transmission of BA.5.2.48 and BA.2 in hamsters. Viral titers of the lungs and turbinates (Tb) of the inoculated donor and the acceptor hamsters were indicated in red and black, respectively. WT hamsters were intranasally inoculated with BA.2, DY.1.1 or DY.2; K18-hACE2 hamsters were intranasally inoculated with BA.2, DY.1.1, DY.2 or WT. Five hamsters per group were used to measure the respective parameters (C, D, F, G and H). Four hamsters per group were euthanized at 3 DPI and used for data collection (E and I). The data (in C, D, F, G and H) of the mock, BA.2, DY.1.1, DY.2 and WT groups are shown in gray, red, blue, orange and green, respectively (as demonstrated in C and F). **(B)** After challenge, the WT hamsters survived, while the K18-hACE2 hamsters died. **(C/G)** Body weights of infected WT (C) or K18-hACE2 (G) hamsters. Significant differences between the mock group and each infected group are revealed above the lines using asterisks in colors corresponding to the respective infected group. The significant differences between DY.1.1 and DY.2 are shown with black asterisks above the lines. **(D/H)**. Relative viral RNA loads in the nasal lavages of infected WT (D) or K18-hACE2 (H) hamsters and baseline viral loads are indicated by dotted gray lines. **(E/I)** Viral titers in the lungs (dark red) or turbinates (cray) of infected WT (E) or K18-hACE2 (I) hamsters. **(F)** Percentage survival of infected K18-hACE2 hamsters. **(J)** H&E staining of the lungs of infected K18-hACE2 hamsters. The selected areas (revealed by a black rectangle in each major panel) are enlarged and shown in inset panels; scale bars: 500 μm (major panels) and 100 μm (inset panels). **(K)** Distribution of virus in the lungs of infected K18-hACE2 hamsters; scale bars: 200 μm.

We then investigated the replication and pathogenicity of DY.1.1 and DY.2 by using more susceptible K18-hACE2 transgenic hamsters. Both WT- and Omicron-infected transgenic hamsters exhibited high mortality (Fig. 3B). The earliest lethality was demonstrated in hamsters infected with the WT virus at 4 DPI (Fig. 3F). Thereafter at 5 DPI, infection of BA.2, DY.1.1 and DY.2 respectively induced 20%, 60% and 80% mortality, respectively, while all hamsters infected with the WT virus died. Significant body weight loss induced by WT infection was revealed as early as 2 DPI (Fig. 3G). Interestingly, the initial state of the three Omicron infections induced a significantly higher increase in body weight than the mock infection at 1 to 3 DPIs, although weight loss was ultimately demonstrated at 5 DPI following BA.2 infection. For all groups, the viral loads in the nasal lavages were roughly equivalent and remained detectable before lethality occurred (Fig. 3H). At 3 DPI, the viral titers in both the lungs and turbinates were higher in the WT group than in the BA.2 and DY.1.1 groups, while no significant difference was demonstrated between DY.1.1 and DY.2 (Fig. 3I). To evaluate the pathological features, we performed H&E staining of the lungs of the hamsters. At 3 DPI, the lungs of the DY.1.1 group exhibited moderate alveolar expansion, exfoliation of bronchial mucosal epithelial cells and severe alveolar wall thickening (Figs. 3J and S5). Moderate alveolar cell necrosis, severe exfoliation of bronchial mucosal epithelial cells and strong infiltration of inflammatory cells in alveolar spaces were demonstrated in the DY.2 group. Similar pathological features with more severe alveolar wall thickening were observed in the WT group. In contrast, BA.2-infected hamsters demonstrated relatively mild pulmonary pathology, which included alveolar wall thickening and inflammatory infiltration in only a limited area. To determine the detailed infection areas in the lungs, anti-nucleocapsid immunohistochemistry was performed. In both the WT and DY.2 groups, widely distributed infection area including the respiratory bronchi epithelium alveolar sacs and the alveoli were clearly observed, and exfoliated nucleocapsid-positive bronchial mucosa formed obstructions in the airways (Figs. 3K and S5). The distribution of DY.1.1 was similar to that of WT and DY.2, while BA.2 tended to infect alveolar sacs and alveoli exclusively. In contrast, less and no airway obstruction was observed for DY.1.1 and BA.2 infection, respectively. Thus both DY.1.1 and DY.2 revealed higher pathogenicity than BA.2, and the pathogenicity of all Omicron variants were lower than WT.

K18-hACE2 transgenic mice, another commonly used susceptible animal model(9, 18), were also used to study the differences in in vivo infection among the three Omicron strains. DY.2 infection of hACE2-expressing mice resulted in 50% mortality at 7 DPI, while all mice infected with BA.2 and DY.1.1 survived (Fig. 4A). All three Omicron infections led to conspicuous and almost continuous body weight loss from 4 to 10 DPI, of which DY.2 infection led to the highest average weight loss (Fig. 4B). However, no significant differences in weight loss were demonstrated among DY.1.1, DY.2 and BA.2 groups. Except for some DY.2-infected individuals, the viral loads in the throat swabs of each Omicron-infected group continued to decrease over 2 to 8 DPI and disappeared at 6 DPI (Fig. 4C). Average viral loads in the DY.1.1 oral swabs were the highest at 2 and 4 DPI. However, in both the lungs and turbinates, there were no significant differences in the viral titers among the three groups (Fig. 4D). Highly consistent with the results preformed in K18-hACE2 hamsters, DY.2 also revealed higher pathogenicity than BA.2 and DY.1.1.

**Fig. 4.**
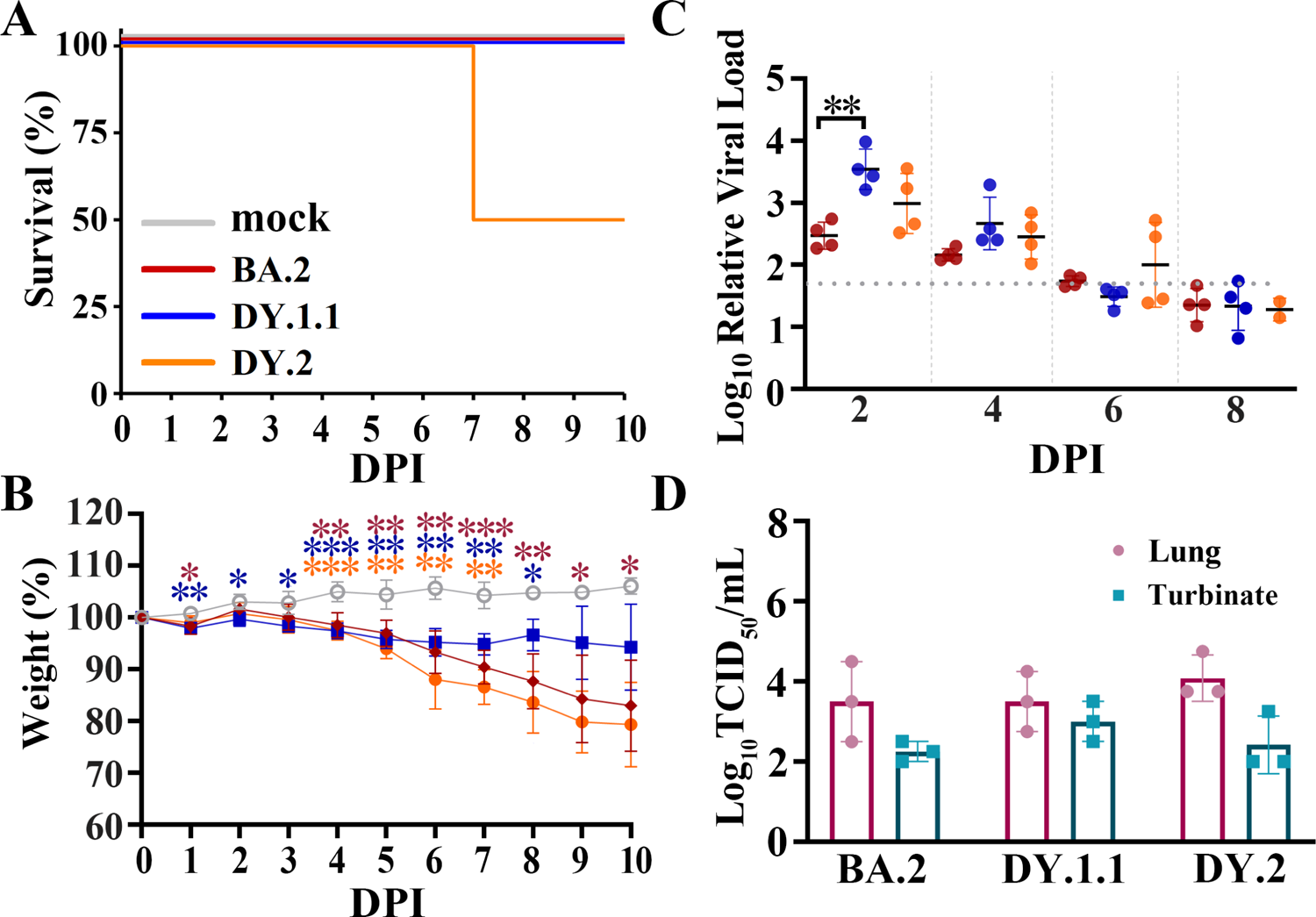
In vivo virological characteristics of BA.5.2.48 in K18-hACE2 mice. Mice were intranasally inoculated with BA.2, DY.1.1 or DY.2, and the corresponding results (A, B and C) are shown in red, blue or orange, respectively (as shown in A). **(A-C)** Four mice per group were used to measure the percentage of survival (A), body weight (B) and relative viral RNA loads in oral swabs (C). Significant differences in Fig. 4B were revealed as in Fig. 3B. **(D)** Four mice per group were euthanized at 3 DPI, after which the viral titers in the lungs (dark red) and turbinates (cray) were measured.

### Competitive replicative fitness of DY.1.1 and DY.2

To further compare the competitive replicative fitness of DY.1.1 and DY.2, hamsters were inoculated with a mixture containing equal amounts of DY.1.1 and DY.2, for which the viral load ratio of DY.1.1 to DY.2 was approximately 0.6. A weak fitness of DY.1.1 over DY.2 was revealed in turbinates of 4/6 hamsters (#1, #2, #3 and #6) at 3 DPI which was invisible in other hamsters (#4 and #5) (Fig. 5A). Nevertheless, divergent results were demonstrated in the lungs, of which a clear dominance of DY.2 was observed in the lungs of one hamster (#5) and the fitness of DY.1.1 was clear in two hamsters (#2 and #6). In contrast to the results at 3 DPI, a mild but stable fitness of DY.1.1 was indicated in both the turbinates and lungs at 5 DPI, except in the turbinate of one hamster (#12). In the nasal lavage samples from all five hamster, DY.1.1 was found to have a slight but stable replicative fitness from 2 to 8 DPI, although DY.2 was predominant at 8 DPI in one hamster (#14).

**Fig. 5.**
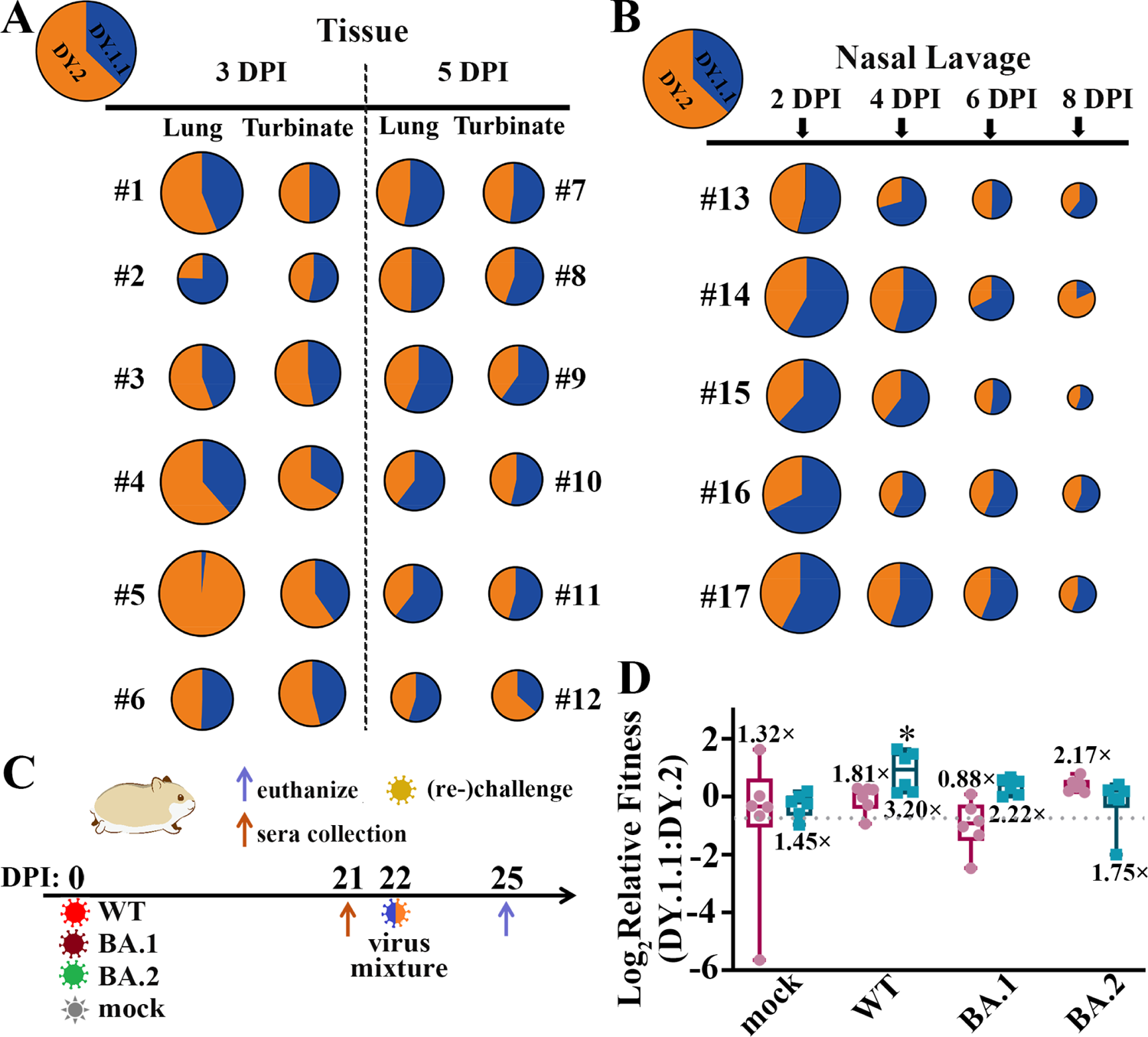
In vivo fitness of the BA.5.2.48 subvariants. **(A)** Competitive fitness of DY.1.1 (blue) and DY.2 (orange) in the lungs and turbinates of coinfected hamsters at 3 DPI (numbered #1 to #6) and 5 DPI (numbered #7 to #12); the proportions of DY.1.1 and DY.2 viral RNA in the inoculation mixture are indicated in the top left corner. (B) Fitness in nasal lavage fluid from coinfected hamsters (numbered #13 to #17) collected at 2, 4, 6 and 8 DPI. **(C)** One day after serum collection at 21 DPI, the (mock-) convalescent hamsters were rechallenged with the DY.1.1 and DY.2 mixture, after which tissue samples were collected three days after rechallenge. **(D)** Relative fitness in the lungs (dark red) or turbinates (cyan) of DY.1.1 compared to DY.2 in mock-, WT-, BA.1- and BA.2-convalesced hamsters. The ratio of DY.1.1 to DY.2 in inoculation is indicated by a dotted gray line (referred to as the inoculation ratio). The ratios of sample ratios to the inoculation ratio were used to determine the relative fitness. Median fitness was demonstrated by lines in boxes above (for fitness in the lungs) or below (for fitness in the turbinates). Significant differences between the convalescent group and the mock group were analyzed.

The results of competitive replication indicated that DY.1.1 slightly outcompete DY.2 in the upper airways of naïve hamsters. We then evaluated the fitness of these strains in hamsters with preexisting immunity. Hamsters were inoculated with WT, BA.1 and BA.2 or were mock infected (naïve). After 21 days, naïve and convalesced animals were (re)challenged with a mixture containing equal amounts of DY.1.1 and DY.2 (Fig. 5C). Compared to the DY.1.1-to-DY.2 ratio in the inoculation (0.6), the ratios in the turbinates of the naïve and WT-, BA.1- and BA.2-convalesced groups were 1.45-, 3.20-, 2.22- and 1.75-fold higher, respectively (Fig. 5D). Compared to those in the naïve group, DY.1.1 in the WT-convalesced group exhibited significant dominance against DY.2 exclusively, indicating that the ability of DY.1.1 to escape from WT serum was greater than that of DY.2. Unlike the convergent DY.1.1-to-DY.2 ratios in turbinates of the naïve group, the divergent fitness of DY.1.1 in the lungs was 1.32 times higher than that in the inoculation. Furthermore, the DY.1.1 fitness levels were 1.81, 0.88 and 2.17 times higher in the lungs of the WT-, BA.1- and BA.2-convalesced groups, respectively; unfortunately, none of these results were significantly different from those of the naïve group. To compare the degree of antibody escape between DY.1.1 and DY.2, the neutralization of WT-convalesced serum against the selected Omicron strains was investigated. However, all the authentic Omicron strains showed striking immune escape from WT infection, and no significant differences were detected between DY.1.1 and DY.2 (Fig. S6).

## Discussion

Mutations in the SARS-CoV-2 spike protein are commonly considered to be highly related to polymorphisms in virological characteristics(9, 10, 16, 19). From former VOCs to Omicron, SARS-CoV-2 evolution has been associated with either enhanced transmission or enhanced humoral immunity evasion(20–23), the former of which confers higher fitness in the upper airways. Within large populations with many infections, the rapid spread of BA.5.2.48 after the end of the dynamic zero-COVID-19 policy in China promoted the generation of multiple subvariants. In contrast to previous virological comparisons between BA.2 and BA.5(3), as offspring of BA.5, DY.1 and especially DY.2, exhibit stronger pathogenicity than BA.2 in K18-hACE2 rodents without enhancement of spike fusogenicity, which suggests that functional alterations in other genes (such as ORF1ab and nucleocapsid in BA.5.2.48 subvariants) also contribute to polymorphisms in virological characteristics. However, spike mutations are also partially associated with BA.5.2.48 characteristics. The fact that DY.1.1 outcompetes DY.2 in the turbinate of hamsters and the high viral loads of DY.1.1 in nasal lavage of rodents suggest its fitness in upper airways, which is positively correlated with the enhanced processing and increased hACE2 binding affinity of the DY.1.1 spike protein. Given that the Chinese population had a high rate of COVID-19 vaccination (anti-WT-spike) before BA.5.2.48 broke out, fitness in WT-convalescent upper airways and putative stronger immune evasion are hypothesized to facilitate the transmission of DY.1.1 (and DY.1). Taking the extremely low prevalence of SARS-CoV-2 in China before the end of zero-COVID-19 policy, our study revealed the putative regulation of SARS-CoV-2 evolution in a relative closed circumstance, and further study could be performed based on our results. What is more, distinct from previous results that Omicron infection is not lethal to the transgenic K18-hACE2 mice and hamsters(3, 18), the H11-K18-hACE2 rodents we used, especially H11-K18-hACE2 hamsters here used which successfully revealing pathogenicity of different Omicron strains, are much more sensitive to Omicron infection and can serve as a better Omicron model.

## Materials and Methods

### Plasmids, PCR and qRT-PCR

The sequences encoding WT, BA.2 and BA.5 spike proteins without the C-terminal 19 amino acids (spike-D19) were synthesized and subcloned into a pcDNA3 plasmid (pcDNA3-spike). A codon-optimized BA.5 spike ectodomain with a furin cleavage site mutation (_682_RRAR_685_ mutated to _682_GSAS_685_), a “HexaPro” modification(24) and a 6×His-StrepⅡ tag was synthesized and subcloned into a pcDNA3 plasmid (pcDNA3-spike-HexaPro). The plasmids expressing the BA.5 spike with the A570S mutation or the A570S/K147E double mutation were obtained by site-directed mutagenesis using a QuikChange II Site-Directed Mutagenesis Kit (Agilent). The dual split protein (DSP) plasmids pDSP1-7 and pDSP8-11 encoding the split Renilla luciferase and GFP genes were synthesized following previous methods(25) and subcloned into the pcDNA3 plasmid. Viral RNA extraction was performed using a QIAamp Viral RNA Mini Kit (Qiagen). The first-strand cDNA synthesis used for reverse transcription (RT), RT-PCR used to amplify specific spike fragments, and qRT-PCR used to determine the relative genome copy number (or viral load) of SARS-CoV-2 were as previously described(26). All primers used are listed in Table S1. To distinguish replicating viral genomes in infected samples from those in aerosol-contaminated samples, the average viral load in samples collected from uninfected animals (mock groups) was used as a detection baseline in qRT-PCR experiments.

### Cells, transfection, protein purification and SPR

Vero E6 cells, HeLa^hACE2+^ cells, HEK-293T cells, Calu-3 cells and HEK-293F cells were cultured and transfected using either Lipofectamine 3000 (Thermo) or polyethylenimine (Polysciences) as previously described(26, 27). HEK-293F cells were transfected with pcDNA3-spike-HexaPro for five days. Spike protein purification was performed following a previous publication(24) with several modifications. Spike proteins in culture medium were purified with gravity flow columns using Strep-Tactin XT (IBA Life Sciences), concentrated to 0.5 mg/ml with ultrafiltration tubes (Millipore) and stored at −80 ℃ before use. The purified proteins were analyzed on 6% SDS-PAGE gels with Coomassie blue staining. The SPR experiments were performed using a Biacore T200 (Cytiva). All assays were performed with 1×HEPES running buffer (10 mM HEPES, 150 mM NaCl, 3 mM EDTA, and 0.005% Tween-20, pH 7.4) at 25 ℃. To determine the binding kinetics between the spike and hACE2 proteins, a Protein A sensor chip (Cytiva) was used. The hACE2 protein with an Fc tag (Acro Biosystems) was immobilized onto the sample flow cell of the sensor chip. The reference flow cell was left blank. Each spike protein was injected over the two flow cells at a range of eight concentrations prepared by serial twofold dilutions (from 3.906 nM to 500 nM) at a flow rate of 30 μl/min using a single-cycle kinetics program. All the data were fitted to a 1:1 binding model using Biacore T200 Evaluation Software 3.1.

### Virus and neutralization

The three SARS-CoV-2 Omicron BA.5.2.48 subvariants used in this study included a DY.1.1 isolate (hCoV-19/Jilin/JSY-CC3/2022, GISAID Accession No. EPI_ISL_18233948), which contains the extra mutations C11750T (NSP6_L260F) and C13166T (NSP10_H48Y); a DY.2 isolate, DY.2-CC1 (hCoV-19/Jilin/JSY-CC1/2023; GISAID Accession No. EPI_ISL_18233990), which contains the extra mutations T2390C (NSP2_F529L) and T11958C (NSP7_I39T); and a DY.2 isolate, DY.2-CC2 (hCoV-19/Jilin/JSY-CC2/2023, GISAID Accession No. EPI_ISL_18233989), which contains the extra mutations G1727T (NSP2_V308L) and C11750T (NSP6_L260F). Other viruses included a WT isolate (IME-BJ05-2020, GenBank Accession No. MT291835), an Omicron BA.1 isolate (hCoV-19/Tianjin/JSY-CC4/2021, GISAID Accession No. EPI_ISL_18435547) and a BA.2 isolate (hCoV-19/Jilin/JSY-CC5/2022, GISAID Accession No. EPI_ISL_18435548). SARS-CoV-2 was passaged in Vero E6 cells as previously described(27). Viral titers were measured using TCID_50_ assays in Vero E6 cells using immunofluorescence as a readout (also see below). An endpoint dilution microplate neutralization assay was performed to measure the neutralization of the authentic virus. Briefly, twofold serial dilutions of sera were made in DMEM supplemented with 10% fetal bovine serum and incubated with 25 TCID_50_ of each indicated virus at 37°C for 1 h. The mixture was then overlaid on 96-well Vero E6 cells. The mean 50% neutralization titer (NT_50_) of the six hamster serum samples was determined using immunofluorescence as a readout.

### Pseudovirus production and infection

The vesicular stomatitis virus (VSV) pseudotyped virus deficient in the G gene (ΔG) bearing a firefly luciferase reporter gene and the VSV-G glycoprotein for infection (VSV-Luciferase-ΔG*G, Brain Case) was passaged in HEK-293T cells transfected with the pMD2.G plasmid. To produce spike-glycoprotein-bearing pseudovirus (VSV-Luciferace-ΔG*Spike), HEK293T cells were infected with VSV-Luciferace-ΔG*G at an MOI of 0.1 after transfection with pcDNA3-spike plasmids (or pcDNA3 as a negative control) and washed twice with DMEM after 2 h according to previous methods(28). Media containing pseudoviruses at 36 HPI was centrifuged at 500 ×g for 10 min, after which the supernatant was stored at −80°C. To measure spike-mediated infection, 2×10^4^ HeLa^hACE2+^ cells (in 96-well culture plates) were mixed with 100 μl of pseudovirus, washed once with DMEM after 24 h and mixed with 100 μl of luciferase substrate (PerkinElmer). Luminescence was detected using an Infinite 200 Pro plate reader (Tecan) as spike-mediated infectivity.

### Cell-cell fusion

Spike-mediated cell-cell fusion was determined following previous publications(9, 25) [9, 23] with several modifications based on the respective split proteins, DSP8-11 and DSP1-7(25), which were expressed in effector and target cells, respectively. In detail, to prepare effector cells, HEK293T cells in 12-well plates were cotransfected with 1 μg of pcDNA3-spike (or pcDNA3 as a negative control) and 1 μg of pDSP8-11. To prepare target cells, HeLa^hACE2+^ cells in 6-well plates were transfected with pDSP1-7 (2.5 μg). A total of 1.3×10^5^ effector cells were transferred into 96-well plates at 24 hours post transfection. After another 24 hours, the target cells were incubated with 1:250 diluted EnduRen live cell substrate (Promega). After detachment, 2.6×10^5^ targeted cells in 100 μl were added to wells containing effector cells. Luminescence was measured at the indicated time points using an Infinite 200 Pro plate reader (Tecan) and used as fusion activity. Cell images were taken using an APX100 microscope (Olympus).

Alternatively, HeLa^hACE2+^ cells in 24-well plates were transfected with 0.2 μg of pEGFPN1 and 0.3 μg of the pcDNA3-spike plasmid (or pcDNA3 as a negative control). Cell images were taken using an APX100 microscope (Olympus).

### Animal experiments

For the airborne transmission study between hamsters, five-week-old male Syrian hamsters (Charles River) were intranasally inoculated with 5000 TCID50 (in 100 μL) of the indicated viruses. Twenty-four hours later, an infected donor hamster and a naive hamster (an acceptor) were cohoused for another twenty-four hours. The donor and receptor were placed in respective small customized transmission cages supporting unrestrained air exchange which were separated by 3 centimeters to prevent direct contact. The donors were placed in front of the isolator unit which provided unidirectional airflow. Tissue samples were collected 3 days after inoculation for the donors or 3 days after initial contact for the acceptors.

Nine 8-week-old male Syrian hamsters (Charles River) or nine 10-week-old female K18-hACE2 Syrian hamsters (H11-K18-hACE2, State Key Laboratory of Reproductive Medicine and Offspring Health, China)(29) per group, including five animals in the body weight subgroup and four animals in the euthanized subgroup, were intranasally inoculated with 5000 TCID_50_ (in 100 μl) of the indicated viruses.

Eight 8-week-old male K18-hACE2 (K18-hACE2-2A-CreERT2) mice (Cyagen Biosciences) per group, including four animals in the body weight subgroup and four animals in the euthanized subgroup, were intranasally inoculated with 2500 TCID_50_ (in 50 μl) of the indicated viruses.

For coinfection studies, DY.1.1 was mixed with DY.2-CC1 at a viral titer ratio of 1:1, and the virus mixture (total 5000 TCID_50_ in 100 µL) was inoculated into WT Syrian hamsters (Charles River), which were either naïve or WT, BA.1-convalescent or BA.2-convalescent. Seventeen 6-week-old male naïve hamsters per group were used for the naïve experiments. For convalescent experiments, 6-week-old hamsters were inoculated with 5000 TCID_50_ (in 100 µL) of WT, BA.1, BA.2 or DMEM (as a naïve group), respectively and 500 μl of blood was taken from the veins of the eyes to collect the serum at 21 DPI. Using RT-PCR to amplify the spike fragment flanking spike_570_ (which contains the _DY.1.1_T23270G_DY.2_ mutation in the SARS-CoV-2 genome) followed by Sanger sequencing, the ratio of DY.1.1 to DY.2 was determined by the peak height ratio of T_DY.1.1_ to G_DY.2_. For all animals, intranasal inoculation, blood collection and euthanization were performed under isoflurane anesthesia.

### Histopathology

Animal lungs were fixed in 4% paraformaldehyde in phosphate-buffered saline (PBS) and processed for paraffin embedding. The paraffin blocks were sliced into 3-µm thick sections and mounted on silane-coated glass slides, which were subsequently subjected to H&E staining for histopathological examination. Panoramic images of the digital slides were taken using a Pannoramic MIDI (3DHISTECH).

### Immunoblotting, immunohistochemistry and immunofluorescence

Virions of SARS-CoV-2 in clear supernatant (10 mL) were inactivated by adding 0.1% paraformaldehyde (final concentration) for 12 h at 4 ℃ and then concentrated by ultracentrifugation at 90000 ×g for 2 h using a P80A rotor (Hettich). The pellets were resuspended in 30 μl of RIPA buffer. The samples were separated via 4-12% SDS-PAGE gels and transferred to PVDF membranes. After blocking with 5% milk, the membranes were blotted with primary antibodies, incubated with horseradish peroxidase-conjugated secondary antibodies and visualized with chemiluminescent reagent as previously described [24]. For immunohistochemistry, tissue sections were processed for immunohistochemistry with a primary antibody and horseradish peroxidase-conjugated secondary antibody and finally stained with a DAB substrate kit (Solarbio). Panoramic images of the digital slides were taken using an SQS-40R slide scan system (Shengqiang Technology). Immunofluorescence was performed as previously described [24]. The antibodies used are listed in Table S2.

### Statistical analysis

All data are presented as means ± SD. Comparisons were performed using student’s test (for repeats no more than 4) or one-way ANOVA. Significance was defined as P<0.05 (*), P<0.01 (**) or P<0.001 (***) and was indicated either in the figures or mentioned separately in the texts. The number of repeats is specified in individual panels using discrete points. All presented data are biological replicates.

### Ethics and biosecurity

All animal experiments were approved by the Animal Care and Use Committee of the Changchun Veterinary Research Institute (approval number: AMMS-11-2023-032). All experiments involving infectious SARS-CoV-2 were performed at the Animal Biosafety Level 3 Laboratories of the Changchun Veterinary Research Institute, Chinese Academy of Agricultural Sciences.

## Supplementary Material

Figs S1 to S6 and Tables S1 to S3 are obtained from Supplementary Materials.

## Contributors

W.W., Q.J., R.L., P.Z, T.L, H.Z. T.W., X.W., and H.X. performed all the virus-related experiments. W.Z., Q.C., Y.G., and Y.L. generated the H11-K18-hACE2 hamsters. X.W. designed the study and wrote the first draft. Y.G., J.L., F.Y., and X.X. revised the first draft and approved the submitted version.

## Acknowledgments

This work is supported by the National Key Research and Development Program of China (2023YFC0871100 to Y.G., 2021YFC2302405 to X.W., 2021YFF0702500 to J.L. and 2023YFC2605500 to Q.C.). This work is also supported by the Natural Science Foundation of Jiangsu Province (BE2019730) and Medical Research Project of Jiangsu Provincial Health Commission (Z2023003) to W.Z.

## Data sharing statement

The original data that support the findings of this study are available from the corresponding author upon reasonable request.

